# Bacterial adaptation by a transposition burst of an invading IS element

**DOI:** 10.1101/2020.12.10.420380

**Authors:** Scott R. Miller, Heidi E. Abresch, Nikea J. Ulrich, Emiko B. Sano, Andrew H. Demaree, Arkadiy I. Garber

## Abstract

The impact of transposable elements on host fitness range from highly deleterious to beneficial, but their general importance for adaptive evolution remains debated. Here, we investigated whether IS elements are a major source of beneficial mutations during 400 generations of laboratory evolution of the cyanobacterium *Acaryochloris marina* strain CCMEE 5410, which has experienced a recent or on-going IS element expansion. The dynamics of adaptive evolution were highly repeatable among eight independent experimental populations and included beneficial mutations related to exopolysaccharide production and inorganic carbon concentrating mechanisms for photosynthetic carbon fixation. Most detected mutations were IS transposition events, but, surprisingly, the majority of these involved the copy-and-paste activity of only a single copy of an unclassified element (ISAm1) that has recently invaded the genome of *A. marina* strain CCMEE 5410. Our study reveals that the activity of a single transposase can fuel adaptation for at least several hundred generations.

**Impact statement:** A single transposable element can fuel adaptation to a novel environment for hundreds of generations without an apparent accumulation of a deleterious mutational load.

## Introduction

The role of transposable elements (TEs) in adaptation has been long debated. These mobile DNA sequences may confer a continuum of phenotypic effects on their hosts [1] but have often been considered to solely be genetic parasites [2,3] with largely deleterious consequences for host fitness [4]. These include the disruption of gene regulation or function following transposition to a new location in the genome, large-scale genomic rearrangements resulting from ectopic recombination, and the generation of double-strand DNA breaks (reviewed by [5]). More recently, however, investigations of insertion sequence (IS) elements – the simplest TEs, found in bacteria and archaea, which consist only of a transposase gene(s) encoding the mobilization machinery [6] – have concluded that a neutral model can explain observed patterns of IS distribution and abundance in bacterial genomes [7,8]. Still, it is well-known that IS elements can sometimes also be beneficial for the host through selectively favored null mutations, modified expression of adjacent genes, or large rearrangements [9–13].

Bacteria and archaea exhibit extensive natural variation in IS element number; most bacterial genomes contain no or few (< 10) elements, while others have hundreds [14–16]. There is also great variation within and between bacterial species in transposition activity and IS-mediated ectopic recombination rates [17]. We therefore expect that the relative importance of transposition for adaptive evolution in bacteria compared with other mutational mechanisms may scale with IS element copy number and activity. IS elements are predicted to contribute little to adaptive evolution when they are rare [18], but they can play a substantial role when moderately abundant. For example, during the initial stages (≤ 500 generations) of adaptation in *E. coli*, transposition or other structural variation involving IS elements (e.g., ectopic recombination) accounted for more than half of beneficial mutations for *E. coli* K12MG1655 (which has 44 TEs) evolved in the mouse gut [19] and for ~25% of genetic diversity in the Lenski long-term evolution experiment [20]. Here, we investigated whether IS elements are the predominant source of beneficial mutations during laboratory evolution of the cyanobacterium *Acaryochloris marina* strain CCMEE 5410 [21,22], which has hundreds of IS elements in its genome [23]. We report that most selectively favored mutations involved the transposition of only a single IS element.

## Results

### Recent IS transposition burst in *Acaryochloris marina* strain CCMEE 5410

Strains of *A. marina* are unique in the production of Chlorophyll *d* as the primary photosynthetic pigment and have large genomes for bacteria, due in part to their high copy number of IS elements [23,24]. We compared IS element copy number in the genomes of *A. marina* strains MBIC11017 [24], CCMEE 5410 [23] and S15 (an epiphyte of the red alga *Pikea pinnata* isolated in 2016 from Shelter Cove, CA), together with the outgroup strain *Cyanothece* strain PCC 7425 (Figure 1A). For this analysis, we used an improved assembly for *A. marina* strain CCMEE 5410 (NCBI BioProject ID PRJNA16707; 23 contigs, N50 = 4,516,345) and genome data acquired for strain S15 (NCBI BioProject ID PRJNA649288; 7 contigs, N50 = 5,881,945). The CCMEE 5410 genome has a much greater number of IS elements compared with the other genomes (Figure 1B). These differences cannot be explained by differences in genome size, which are comparable for *A. marina* genomes (8.09 Mbp for CCMEE 5410 versus 8.36 Mbp for MBIC11017 and 7.11 Mbp for S15). IS element transposase genes account for ~8% of protein-coding genes in the CCMEE 5410 genome and include high element copy numbers for IS families that are either absent from or have a low copy number in the genome of sister taxon strain MBIC110017 (e.g., ISAs1; Figure 1 – figure supplement 1).

**Figure 1.**
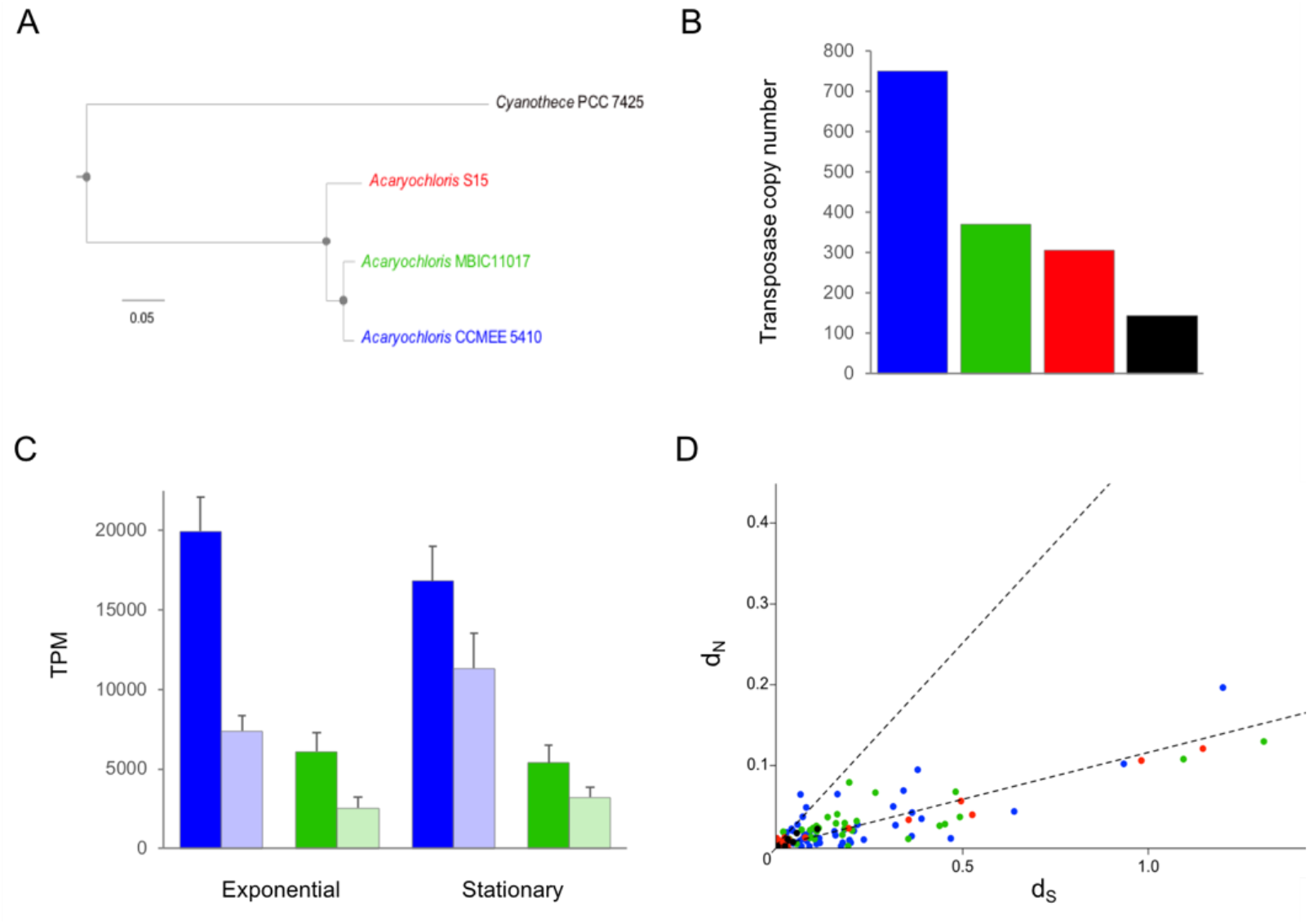
IS element expansion in *Acaryochloris marina* strain CCMEE 5410. **(A)** Maximum likelihood amino acid phylogeny of *A. marina* strains CCMEE 5410, MBIC11017 and S15, outgroup-rooted with *Cyanothece* strain PCC 7425. The tree was reconstructed from a concatenated alignment of 1468 orthologous proteins using a JTT+F+R5 substitution model. All nodes had 100% bootstrap support for 1,000 bootstrap replicates (indicated by closed circles). Scale bar is in units of expected number of amino acid substitutions per site. **(B)** Genome-wide number of transposase genes for each of the four strains. Color coding as in (A). **(C)** Exponential growth and stationary phase expression (transcripts per kilobase million) of sense (dark shading) and anti-sense (light shading) transposase gene transcripts for *A. marina* strains CCMEE 5410 and MBIC11017. Error bars are standard deviations. Color coding as in (A). **(D)** Scatter plot of nonsynonymous (d*N*) versus synonymous (d*S*) nucleotide divergence between recent, non-identical transposase gene duplicate pairs for the four strains. For regression analyses, data were pooled from the four strains (excluding 11 duplicate pairs in strain CCMEE 5410 with d*N*/d*S* > 0.3, for which a separate regression line was estimated; see main text). Least-squares regression slopes for the individual strains were as follows: CCMEE 5410 (d*N*/d*S* = 0.12; adjusted *R*^2^ = 0.65; *N* = 60 duplicated copy pairs); MBIC11017 (d*N*/d*S* = 0.13; adjusted *R*^2^ = 0.88; *N* = 63); S15 (d*N*/d*S* = 0.10; adjusted *R*^2^ = 0.97; *N* = 32); *Cyanothece* PCC 7425 (d*N*/d*S* = 0.09; adjusted *R*^2^ = 1.0; *N* = 10). Color coding as in (A).

IS element expression comprised a disproportionately greater fraction of the CCMEE 5410 transcriptome compared with MBIC11017 than would be expected given the two-fold difference in element number between the genomes (Figure 1C). This was the case for both sense and antisense transcripts and implies that CCMEE 5410 may have less regulatory control over the transcription of IS elements than MBIC11017. In CCMEE 5410, approximately 2% of sense transcripts were derived from IS elements during both exponential growth (mean ± SD = 2.0% ± 0.22%) and stationary phase (1.7% ± 0.22%), respectively. Because unnecessary gene expression is costly [25,26], we consequently expect IS expression to be a greater metabolic burden for CCMEE 5410.

Most IS elements in the CCMEE 5410 genome appear to be of recent origin based on the low levels of synonymous nucleotide divergence (d*S*) among duplicated gene copies within IS families (Figure 1D). Many of these are pseudogenes that have been inactivated by small deletions or insertions but have not yet been purged from the genome; the CCMEE 5410 genome has a much higher percentage of these frameshifted IS remnants (33%) than either the MBIC11017 (14%) or S15 genomes (8%; Supplementary file 1).

We used the ratio of nonsynonymous to synonymous nucleotide divergence (d*N* /d*S*) between recently duplicated full-length transposase gene copies as a measure of the strength of selection on IS elements. This indicated that IS elements are generally under similarly strong purifying selection in these genomes (d*N*/d*S* = 0.12; adjusted R^2^ = 0.88; Figure 1D; p = 0.45 for the *F* test that there is a difference among strains). This level of selective constraint is similar to what has been observed for other retained gene duplicates in *A. marina* [23]. These conserved IS elements may potentially have been domesticated for host function [27]. An alternative explanation for such a high degree of functional constraint on most IS elements is that there is selection against inactivating mutations that result in mis-folded proteins [28]. However, the CCMEE 5410 genome also harbors a small number of recently duplicated transposase gene copies that are experiencing lower selective constraint (Figure 1D; d*N*/d*S* = 0.45; *R^2^* = 0.67; *N* = 11 duplicate pairs; p = 0.001 for an *F* test comparing this high d*N* / d*S* class with all other d*N* / d*S* pairs from all strains). These less constrained transposases were also significantly more highly expressed than more conserved IS elements under both exponential growth and lag phase conditions (Figure 1 – figure supplement 2).

Together, the above observations suggest that the high IS copy number in the *A. marina* CCMEE 5410 genome is the product of a recent or on-going expansion of IS elements from several IS families since it last shared a common ancestor with MBIC11017. This may be a consequence of a reduction in the ability of selection to purge these genes from the CCMEE 5410 genome due to a lower historical effective population size compared with other *Acaryochloris*, similar to the increased number of IS elements (and pseudogenes) observed in the genomes of many obligate bacterial endosymbionts following a history of bottlenecks [29,30].

### Major role for the transposition of a single IS element during adaptive laboratory evolution

The *A. marina* CCMEE 5410 genome provides an opportunity to address the consequences of a high TE load for evolution. To evaluate the relative contribution of IS activity to CCMEE 5410 evolution compared with other mutations, we conducted a laboratory evolution experiment with eight replicate populations (A-H) descended from an ancestral population stock culture (see Materials and Methods). Experimental conditions were identical to the ancestral maintenance conditions, with the exception of the culture volume (150 mL in 250 mL flasks during the experiment, compared with 50 mL in 125 mL flasks for routine maintenance). Population growth was biphasic under these batch culture conditions (Supplementary file 2): after a lag, a period of exponential growth was followed by slower linear growth. Every three weeks (approximately seven generations), 1 mL of culture (~450,000 cells) was transferred into fresh medium. The experimental populations were maintained in this way for 400 generations (approximately 40 months). By the end of the experiment, cells from the evolved populations were ~15% smaller in diameter than the ancestor (mean ± SE of 2.0 μm ± 0.04 versus 2.3 μm ± 0.11). In aggregate, the evolved populations grew ~15% faster during the exponential phase compared with the ancestor (Figure 2A; *t* = 2.59; df = 23; *P* = 0.016). By contrast, no differentiation between the evolved populations or the ancestor was observed during other phases of the growth cycle or for cell yield.

**Figure 2.**
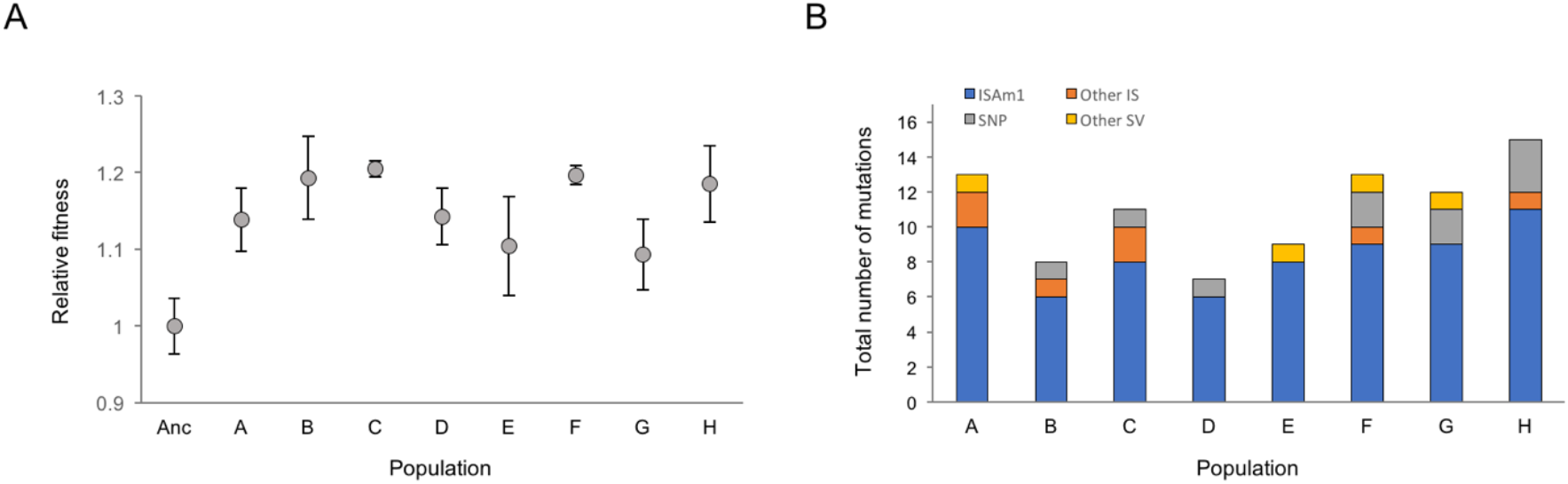
Fitness data and distribution of mutations for laboratory evolved populations of *A. marina* strain CCMEE 5410. **(A)** Relative exponential growth rates of experimental populations after 400 generations of laboratory evolution, compared with the ancestral population (Anc). In this experiment, a relative fitness of 1 corresponded to a population growth rate of 0.26 doublings per day. Error bars are standard errors for biological triplicate cultures. **(B)** Distribution of mutations detected in the populations during the course of the experiment shows the massive contribution of ISAm1 insertions.

To identify the mutations responsible for the observed increase in fitness, every 100 generations we Illumina-sequenced DNA isolated from each population to greater than ~30X coverage (Supplementary file 3; NCBI BioProject ID ###########.). Most detected mutations were IS transposition events (Figure 2B; 75-92% of mutations within each population). As predicted, this is a greater fraction than what has been previously observed in laboratory evolution experiments with *E. coli*, which has fewer IS elements [19,20]. However, the overall evolutionary rate of the *Acaryochloris* populations was similar to what has been observed for *E. coli* over a comparable number of generations [19,20]; at the end of the experiment, individual CCMEE 5410 cells were expected to have ~1-3 mutations. We detected 39 distinct insertion alleles that were not found in the ancestor. Many of these were observed in multiple populations (Figure 2 – source data 1) and are probably the result of convergent evolution (see below). Nearly two-thirds of these insertions (*N* = 25) were in coding regions and are therefore likely null mutations, in accord with the idea that loss-of-function mutations can play an important role in adaptation [12].

Remarkably, the overwhelming majority of these transposition events (≥ 80%) involved an unclassified IS element (ISAm1) that consists of a single DDE transposase gene with a 14-bp inverted repeat (Figure 2B; Figure 2 – source data 1). The direct repeats flanking the detected ISAm1 insertion sites have an average GC content of 27% (Figure 2 – source data 1), suggesting a bias toward AT-rich sites (genome-wide GC content is 47.5% in coding regions versus 41.5% in non-coding regions; Figure 2 – figure supplement 1). ISAm1 appears to have recently invaded the CCMEE 5410 genome, since it is not observed in the other *A. marina* strains. It is, however, homologous to a transposase gene from the cyanobacterium *Moorea* sp. (NCBI accession number NEP53674.1; 68% amino acid identity).

The genome of the CCMEE 5410 ancestor has nine nearly identical ISAm1 copies (Figure 2 – figure supplement 2). However, only one copy (genome coordinates 6:36060-6:37572) is complete; the others appear to be pseudogenes based on one or more premature stop codons resulting from frameshift mutations. Only the complete ISAm1 copy has 100% nucleotide identity with the reconstructed mRNA transcript (Figure 2 – figure supplement 2), suggesting that it (and potentially its descendant copies) is the only transpositionally active copy; the other copies may be nonautonomous but possibly mobilized by this copy. ISAm1 transposition was by a copy-and-paste mechanism, and, at the end of the experiment, the number of ISAm1 copies segregating within populations had increased by 1-5 copies. In the ancestor, ISAm1 was transcribed throughout the batch growth cycle but exhibited highest expression (and highest ratio of sense versus anti-sense transcripts) during lag phase (Supplementary file 4).

### Repeatability of adaptation and the resolution of clonal interference

Drift is expected to be weak compared with selection under our experimental conditions (Ne is > 10^5^ in the evolving populations). Consequently, mutations that rise to a detectable frequency in the population are likely either selectively favored or genetically linked to a beneficial mutation. The observation of identical or parallel mutations in the same target among populations is typically considered to be strong evidence that the locus itself was the target of positive selection. Evolution was highly repeatable among populations and characterized by: (1) the purging of ancestral polymorphism; (2) the subsequent emergence of high frequency ISAm1 insertion alleles in the carbon regulatory operon *sbtAB;* and (3) the resolution of clonal interference among *sbtAB* alleles as additional, often convergent beneficial mutations arose on different genetic backgrounds. We discuss these dynamics in more detail below.

Illumina sequencing of the ancestral population to > ~250X coverage (range: 246-356X; Supplementary file 3) revealed several polymorphisms (Supplementary file 5). These included two derived nonsynonymous SNPs: one in lipoate synthase *lipA* (66% frequency) and the other in *argC* of the arginine biosynthesis pathway (7.5%). Structural variants included a low frequency, 3-bp in-frame insertion in a glycine dehydrogenase gene and two ISAm1 insertion polymorphisms at low (~5%) frequency. All of this ancestral variation was eventually lost in all of the evolved populations, most by generation 100. After 100 generations, we also detected an identical ISAm1 insertion between the urease accessory protein coding genes *ureF* and *ureG* in all populations (Figure 3; Figure 2 – source data 1). This mutation was not detected in the ancestral population and may reflect an insertion hot spot, but we cannot rule out that it was segregating in the ancestral population at very low frequency. This mutation was also lost in all populations later in the experiment (Figure 3).

**Figure 3.**
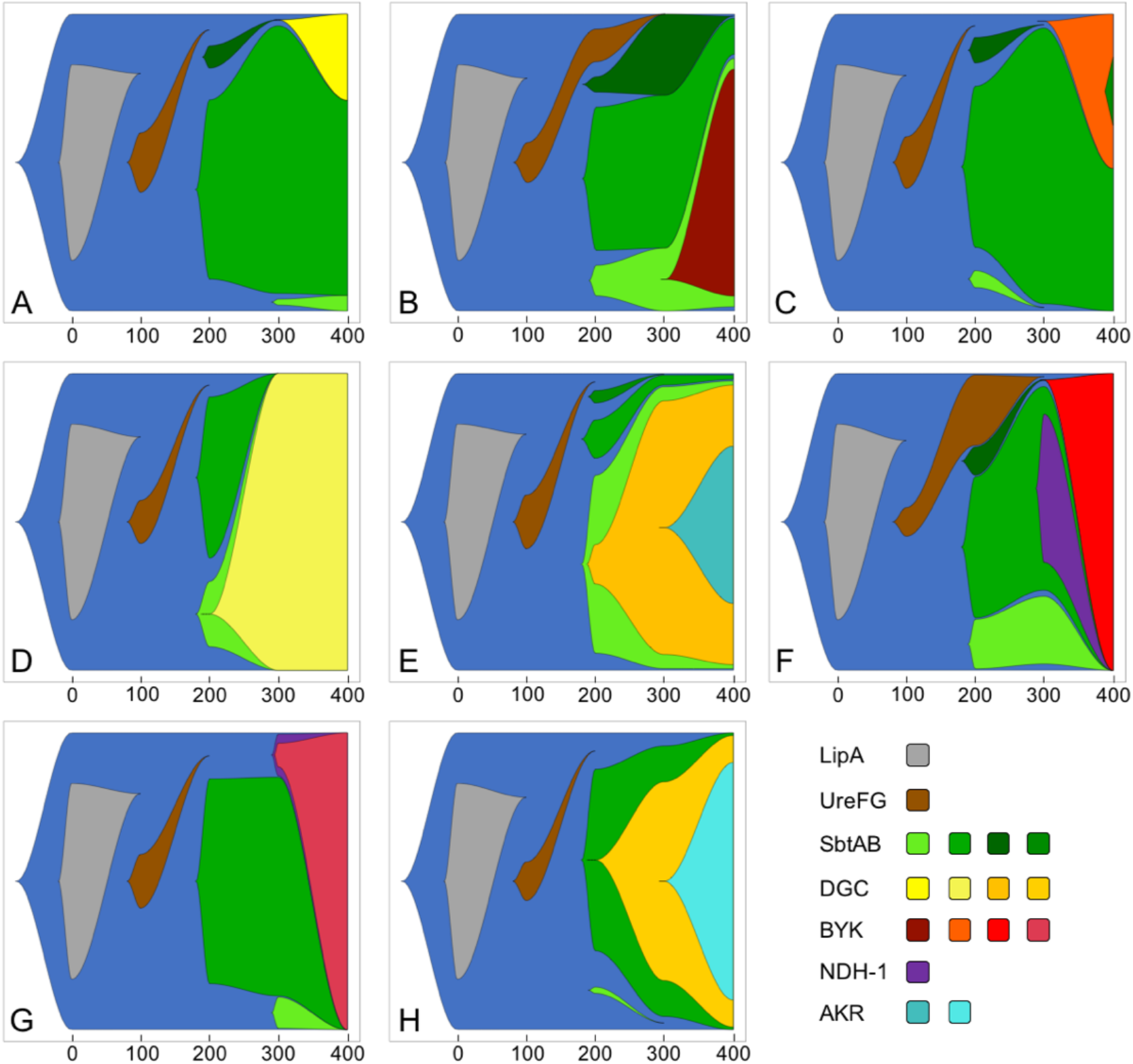
Fish plots of major evolutionary changes during 400 generations of laboratory evolution of the eight populations. The majority of the featured mutations that were detected during the experiment are ISAm1 insertion events. The exceptions are DGC mutations in the A and D populations and BYK mutations in the B, C, and F populations.

By generation 200, between one and three different ISAm1 transposition-mediated alleles were detected in the *sbtAB* operon in all populations (Figure 3). All three alleles were intergenic (a fourth ISAm1 insertion event in *sbtB* emerged in a single population late in the experiment), and two (Sbt-1, Sbt-2) were observed in all populations. For multiple reasons, we believe that these mutations were convergent rather than standing variation. None of the alleles were observed in any population prior to generation 200 (despite sequencing populations to greater than ~250X coverage in generation 100; Supplementary file 3), yet one or more increased rapidly in frequency once detected (Figure 3; Figure 3 – figure supplement 1). This suggests that they were under strong positive selection and would have been detected earlier in the experiment if they had been present in the ancestral population. In addition, we would expect to have observed similar evolutionary trajectories across populations if they were derived from standing variation.

Together, *sbtA* and *sbtB* are involved in cellular acclimation to low carbon. The *sbtA* gene encodes a sodium-dependent, high-affinity bicarbonate transporter that is a part of the cyanobacterial carbon-concentrating mechanism [31], and the *sbtB* product is a P_II_-like cAMP-binding signaling protein involved in sensing cellular inorganic carbon (C_i_) status [32] and regulating SbtA activity. *Synechocystis* PCC 6803 mutants with a *sbtB* deletion are constitutively in a low carbon-adapted state and sensitive to sudden changes in C_i_ supply [32]. In the CCMEE 5410 ancestor, *sbtAB* genes are co-expressed as a single ~1.8 kb bicistronic transcript that is upregulated to 10-fold higher levels during carbon limitation (Supplementary file 6). Whether the *sbtAB* insertions impact C acquisition (e.g., via changes in the stoichiometry of SbtA and SbtB) remains to be determined.

A number of other detected mutations were also associated with C_i_ uptake (Figure 2 – source data 1). These included identical ISAm1 insertions into a *sbtA* homolog (45% amino acid identity to SbtA) that was observed in seven of the populations. We also identified an ISAm1 insertion upstream of the NDH-1MS complex in three populations (Figure 2 – source data 1; Figure 3). NDH-1MS is a cyanobacterial NAD(P)H:Quinone oxidoreductase complex specialized for high affinity CO_2_-uptake under low C_i_ conditions [33]. Similar to what was previously reported for *Synechocystis* PCC 6803 [34], ancestral CCMEE 5410 exhibited increased transcription of these genes in a low C_i_ environment, as were other carbon concentrating mechanism genes (Supplementary file 6). None of these mutations were detected until generation 200 or later, which indicates that they were independently acquired in the individual populations.

The emergence of multiple co-occurring Sbt alleles is expected to produce clonal interference dynamics [35], whereby competition between competing beneficial alleles slows the loss of variation from the population. Still, by the end of the experiment, Sbt diversity was lost in six of the eight populations (1-3 detected alleles versus a maximum of 3-4 alleles), and a single allele had attained high frequency (Figure 3; Figure 3 – figure supplement 1). Four of the five Sbt alleles became dominant in at least one population. This included the ancestral allele, which appeared to be generally selected against, since it was either undetectable or at a low frequency by the end of the experiment in most populations. However, in two populations (C, G; Figure 3), there was a substantial increase in the ancestral allele’s frequency between generations 300-400 as a result of new beneficial mutations that overcame this deleterious genetic background.

The evolutionary outcome of Sbt clonal interference depended upon the genetic background of subsequent beneficial mutations. In three populations, sweeps of a particular Sbt allele (Sbt-1 in the D and E populations, Sbt-2 in H; Figure 3) were associated with mutations either within or upstream of a diguanylate cyclase gene (peg.4655; Figure 4; Figure 2 – source data 1). Mutations at this locus were very common following the emergence of Sbt variation. We observed a total of eight distinct alleles in seven of the populations (Figure 4); the majority of these interrupted the coding region and are therefore expected to be null mutations. Seven of the mutations were due to the transposition of IS elements (five by ISAm1 activity); the other (the D population allele) was a C-to-T mutation resulting in a premature stop codon. Diguanylate cyclases are involved in the production of the secondary messenger molecule cyclic diguanylate, which activates specific effector proteins to impact a number of cellular processes, including biofilm formation and stress responses [36]. Evolutionary changes in cyclic diguanylate signaling have been previously shown to be central to diversification in biofilms of *Pseudomonas aeruginosa* [37]. In CCMEE 5410, peg.4655 is constitutively expressed (Supplementary file 6), and its ortholog in *A. marina* MBIC11017 is upregulated under microoxic conditions [38]. Its effector protein and the downstream consequences of its inactivation remain to be determined.

**Figure 4.**
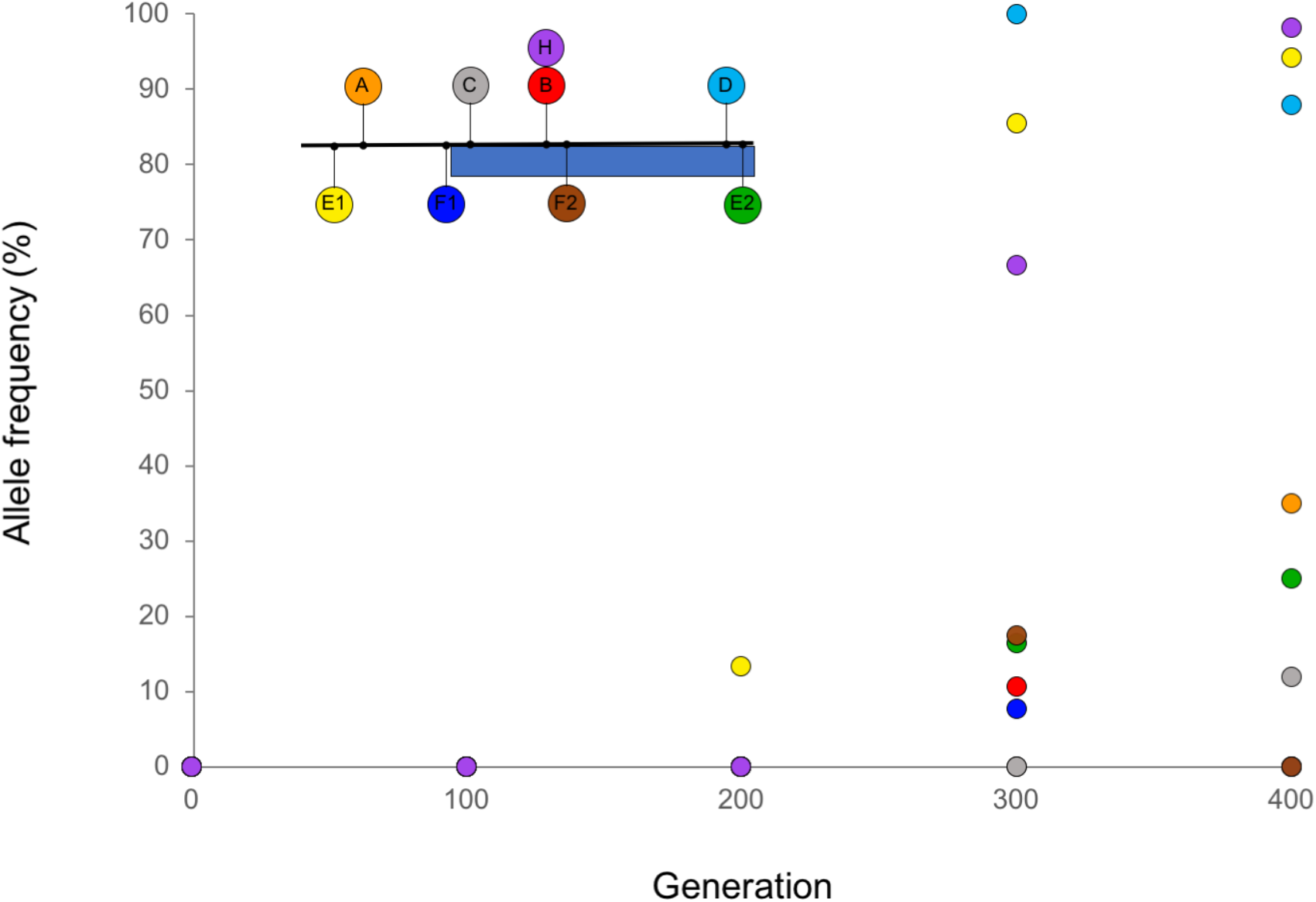
Location and frequencies of mutations detected during laboratory evolution in and near the annotated diguanylate cyclase gene peg.4655. Shown is a 1 kb region of the CCMEE 5410 genome (genome coordinates 0:4457436-0:44578436) including peg.4655 (blue rectangle) and upstream non-coding DNA. All mutations are IS transposition events, with the exception of the D allele, which is a nonsense mutation at amino acid position 207 (Figure 2 – source data 1).

In four populations, by contrast, late-arising mutations in a bacterial tyrosine kinase (BYK) gene (peg.5255) had attained high frequency (53-100%) by the end of the experiment (Figure 3; Figure 2 – source data 1). We detected three nonsynonymous SNPs and two transposition events involving IS families ISAm1 and ISAcma36, respectively. BYKs are signaling proteins that regulate traits such as virulence, stress responses and exopolysaccharide production by both autophosphorylation and substrate phosphorylation of tyrosine residues [39]. The insertions, which are located at sites eight nucleotides apart at the 3’ end of the gene, are expected to disrupt the C-terminal tyrosine cluster autophosphorylation sites of the protein. This could potentially disrupt interactions with its target substrate proteins. This gene also possesses a N-terminal GumC domain, which suggests that it is involved in exopolysaccharide biosynthesis. We predicted that mutations at this locus are associated with the loss of the ability to form biofilm. The results of an adherence assay [38] demonstrated that this was indeed the case. Evolved populations with a BYK mutation produced less biofilm than either evolved populations without a BYK mutation (*F*_1,37_ = 23.6, p < 0.0001) or the ancestral population (*F*_1,19_ = 5.51, p = 0.03). Evolution of a more planktonic lifestyle consequently appears to be advantageous, possibly due to agitation in the selected environment. By contrast, populations lacking a BYK mutation had not diverged from the ancestral value (p = 0.74).

## Discussion

Here, we have shown that a single active copy of a TE can fuel the initial stages of adaptation over hundreds of generations. Copy number of the ISAm1 element expanded during laboratory evolution and was responsible for about 75% of beneficial mutations. This greatly increased the rate of adaptive mutations compared with nucleotide substitutions alone, as has been observed for an IS transposition burst during *E. coli* adaptation to a change in osmolarity [40].

Many of the ISAm1 insertions were in or near genes involved in C_i_ concentration and acquisition (Figure 3; Figure 2 – source data 1). The phenotypic consequences of these mutations will be investigated in detail elsewhere. However, we can identify at least two ways in which C_i_ acquisition may have been under selection during laboratory evolution. First, our experimental treatment imposed a general reduction in the ratio of gas exchange surface area to culture volume compared with the ancestral maintenance conditions. Therefore, environmental C_i_ availability is expected to be lower. C_i_ availability is also expected to fluctuate during the course of a growth cycle, with higher availability during early growth at low cell densities, followed by C-limitation later in the cycle. The nature of selection likely varied temporally as a result.

The predominance of a single TE for adaptation was striking in light of the fact that multiple IS families are actively expressed by *A. marina* CCMEE 5410 (Supplementary file 4). The reasons why we did not observe a more equitable contribution to adaptation from different IS families (including other recently acquired elements that are unlikely to have been domesticated) are not clear. Potentially, insertion site targets are more restricted or saturated for other highly expressed elements. Our study also illustrates the ecological differences among IS elements, which may transpose during different phases of the experimental growth cycle (Supplementary file 2). For example, the ISAm1 element appeared to be particularly transcriptionally active during lag phase, whereas the IS630 family exhibited highest expression during exponential growth (Supplementary file 4). Consequently, the spectrum of IS-mediated mutations available to a bacterium may depend on its current or predominant physiological state [41].

The long term fate of the ISAm1 element is not clear. Simulation studies of both sexual diploid and asexual populations have indicated that an invading TE is more likely to be stably maintained in a genome following an initial transposition burst if its activity is subsequently regulated [42,43]. Otherwise, it is ultimately expected to go extinct, provided that deleterious transpositions are much more common than adaptive insertions. In our experiment, beneficial ISAm1 transposition mutations with a large selective effect were sufficiently frequent to co-occur within a population (Figure 3), corresponding to a strong-selection strong-mutation regime [44]. However, we did not observe any compelling evidence for potentially deleterious ISAm1 transposition mutations hitchhiking to high frequency. Rather, the rare cases of multiple ISAm1 transposition events sweeping together often involved insertions that convergently occurred in multiple populations and were plausibly adaptive. For example, in the G population, there was a rapid sweep of three ISAm1 transposition events between generations 300 and 400 at loci that convergently rose to high frequency in other populations (bacterial tyrosine kinase, coproporphyrinogen III oxidase, and the NDH-1MS complex; Figure 3; Figure 2 – source data 1). Therefore, while beneficial ISAm1 transpositions were frequent enough to compete with each other, the probability of a deleterious transposition event hitchhiking along appears to be low. This suggests that deleterious transposition events may cause strong fitness effects and be effectively purged from the population, preventing the accumulation of a substantial deleterious ISAm1 load that might lead to extinction.

## Materials and Methods

### Strain maintenance

*A. marina* strain stocks were maintained at 30 °C in 125ml Erlenmeyer flasks containing 50 mL of HEPES-buffered (10mM final @ 8.0pH) FeMBG-11 medium (IOBG-11 supplemented with iron(III) monosodium salt; [45]). Cultures were grown with constant shaking (92 rpm) on a VWR Advanced Digital Shaker and illuminated with 25 μmol m^−2^ s^−1^ of cool white fluorescent light on a 12h:12h light:dark cycle.

### Genome data and analysis

Both short-read (Illumina) and long-read (PacBio) genome sequence data were acquired for *A. marina* strains CCMEE 5410 and S15. For CCMEE 5410, cells for Illumina sequencing were obtained directly from the ancestral stock culture used to inoculate the laboratory evolution population cultures (see below). For PacBio sequencing, 1 mL each of the ancestral stock was inoculated into two flasks of FeMBG-11/HEPES and harvested after ~10 generations of growth.

For Illumina sequencing, 120 μl of lysozyme (10mg/mL) were added to a microfuge tube containing approximately 0.1 g of pelleted culture. The tube was next vortexed and incubated at 37 °C for 30 min. Following this, DNA was extracted with the Qiagen DNeasy PowerBiofilm kit according to manufacturer instructions. DNA was Qubit quantified and sent to the University of Pittsburgh Microbial Genome Sequencing Center for library preparation and 151-bp paired-end sequencing on an Illumina NextSeq 500 flow cell.

In addition, high molecular weight DNA was extracted for PacBio sequencing from 100 mL of culture split into two pellets. Each pellet was resuspended in 4.7 mL of TE buffer (pH 8.0). We next added 100 μl of 200 mg/mL lysozyme to each tube and incubated at 37 °C for 45 minutes. Following this, 50 μl of Proteinase K were added, and the tubes were incubated at 55 °C for 1 h. 900 μl of 5M NaCl were then added to each tube, followed by 750 μl of CTAB/NaCl (10 g cetyl trimethylammonium bromide) and 4.09 g NaCl). After incubation at 65 °C for 20 min, cell debris was pelleted at 5,000 x *g* for 10 min at room temperature. The supernatant was transferred to a new tube to which an equal volume of chloroform was next added. The tube was then centrifuged at 5,000 x *g* for 30 min. Following this, the aqueous phase was harvested, and DNA was precipitated with 2X volume of 100% ethanol and then pelleted at 5,000 x *g* for 30 min. 200 μl TE was added to dissolve the pellet, and the solution was transferred to a clean microfuge tube. 200 μl of phenol/chloroform (1:1) was added to the tube, mixed well by repeated inversion, followed by centrifugation for 10 min at 17,000 x *g*. The aqueous layer was then transferred to a clean microfuge tube and extracted with chloroform an additional time as above. DNA was reprecipitated with ethanol as above, and then, after removing the supernatant, resuspended in 50 μl of 3M sodium acetate (pH 5.2). We next added 10 μl of glycogen and 3.5X volume of 100 % ethanol, followed by incubation at −80 °C for 30 min. The sample was then centrifuged at 17,000 x *g* and 4 °C for 15 min. Following this, the supernatant was removed, and the sample was air-dried, resuspended in 10 mM Tris and stored at −80 °C.

Sample quality was assessed with an Agilent Tapestation and by Qubit and Nanodrop. Sequencing was conducted with a PacBio Sequel System at the University of Maryland Institute for Genome Sciences. Genomes for *A. marina* strains CCMEE 5410 and S15 were *de novo* assembled with Canu v1.7 [46], and these assemblies were improved with Pilon [47] using Illumina data. Genome data acquired for this study are available at NCBI BioProject ID PRJNA16707 (CCMEE 5410) and PRJNA649288 (S15).

### Phylogenetic analysis

Orthologous protein-coding genes were identified for the outgroup strain *Cyanothece* PCC7425 (NCBI accession: GCA_000022045.1) and for *A. marina* strains MBIC11017 (GCA_000018105.1), CCMEE 5410 and S15 using OrthoFinder v2.2.7 [48]. A maximum likelihood amino acid phylogeny with 1,000 ultrafast bootstrap replicates [49] was constructed with IQ-TREE v2.0 [50] using the JTT substitution matrix with empirical amino acid frequencies (+F) and five estimated free rate categories of rate heterogeneity among sites (+R5). The model was selected by the Akaike information criterion (AIC) with ModelFinder [51].

### IS element analyses

Genome-wide estimates of transposase gene copy number were obtained by parsing annotation data with a custom Python script. To identify which transposase genes were related by gene duplication and to measure the amounts of synonymous and nonsynonymous nucleotide divergence between pairs of transposase duplicates, we developed a novel bioinformatics software, ParaHunter, which is freely-available on GitHub: https://github.com/Arkadiy-Garber/ParaHunter. ParaHunter identifies homologs by clustering genes using *mmseqs2* v6.f5a1c [52], based on user-chosen parameters of minimum amino acid identity and coverage. After gene clusters are identified, each cluster is aligned using *Muscle* v3.8.1551 [53]. ParaHunter then uses *codeml* in PAML to generate codon alignments (*pal2nal.pl*) and estimate rates of synonymous (d*S*) and nonsynonymous (d*N*) divergence [54]. The resulting d*N* and d*S* values are then extracted from the output files and d*N*/d*S* values calculated directly from these estimates.

To identify gene duplicates in *Acaryochloris* strains, clustering by *mmseqs* required coverage of at least 50% over the length of the target sequence, with a minimum amino acid identity of at least 50% over the length of the shorter sequence. Genes were annotated by comparison with the annotated genome of *Acaryochloris* MBIC 11017 [24] using *DIAMOND BLASTp* v0.9.24.125 [55]. Annotation data were also used to confirm the accuracy of gene clustering, where all members of each cluster of homologous genes are annotated with the same function.

To estimate the amount of nonsynonymous (d*N*) and synonymous (d*S*) nucleotide divergence between pairs of paralogous IS genes, we ran *codeml* on the codon alignments generated using the *pal2nal.pl* script with the following parameters: runmode = pairwise, seqtype = codons➔AAs, model = empirical, NSsites = 0, icode = universal, fix_kappa = kappa to be estimated, fix_omega = estimate, fix_alpha = fix it to alpha, RateAncestor = 1, Small_Diff = 0.5e-6, fix_blength = random, method = simultaneous. Regression analysis of d*N* and d*S* values was performed in *RStudio* (R Core Team, 2013). Analysis of variance (ANOVA) was performed using the base R function *aov(*). In our analyses, we also used the packages *tidyverse* (https://cran.r-project.org/web/packages/tidyverse/index.html) and *reshape* [56].

Pseudogenes were identified using the Pseudofinder software (https://github.com/filip-husnik/pseudofinder) with default parameters and four cyanobacterial reference genomes (*A. marina* strain S15, *Cyanothece* sp. PCC 7425, *Thermosynechococcus elongatus* BP-1 and *Synechococcus* sp. PCC 6312). Frameshifts and insertions/deletions in identified pseudogenes were determined using a custom Python script to parse alignments of duplicated transposase genes for sequence lengths and alignment gaps that are not multiples of three.

RNASeq read data obtained for *A. marina* strains CCMEE 5410 and MBIC11017 [57; NCBI BioProject ID PRJNA681975] were quality-trimmed using Trimmomatic v0.39 (ILLUMINACLIP:TruSeq3-PE:2:30:10 LEADING:3 TRAILING:3 SLIDINGWINDOW:4:15 MINLEN:36) [58]. Given the heavy load of IS gene duplicates (including nearly-identical duplicates) in the *Acaryochloris* strains MBIC11017 and CCMEE 5410 genomes, we performed read mapping using a custom approach that allowed us to keep accurate track of which reads map ambiguously. To estimate expression of genes present in multiple copies in each genome, we utilized a combination of *bowtie2* and *BLASTn. Bowtie2* v2.3.4.3 (default settings) [59] was used to recruit reads separately to each cluster of paralogous genes. To accurately estimate expression levels from each gene within each cluster, while keeping track of ambiguously-mapping reads, the subset of reads mapping to each gene cluster was then queried against its respective gene cluster using BLASTn v2.9.0+ (qcov_hsp_perc = 100%, perc_identity = 100%) [60]. A custom Python script was then used to process the results and estimate *total* read counts from each gene cluster, as well as unambiguous read counts from each *individual* gene within each cluster. Gene expression from single copy genes was estimated using only *bowtie2* (default settings), and the read count estimates were generated using *htseq-count* v0.11.2 [61]. Gene expression values were generated by normalizing the read count estimates to transcripts per million (TPM) [62]. The TPM values reported for each gene/gene cluster and each time point represent the mean and standard deviation from five replicates. Transcriptomes from each time point were assembled using the default settings in *Trinity* v2.8.4 software [63]. All custom Python scripts used here are available in Supplementary file 7.

### Laboratory evolution experiment

We established eight replicate populations (A-H) descended from an ancestral stock culture. Experimental populations were initiated by inoculating 1 mL each from the ancestral stock into 250 mL longneck flasks containing 150 mL of FeMBG-11/HEPES (10mM final, pH 8.0) medium. Experimental medium, temperature and light regime were identical to the ancestral maintenance conditions. Every three weeks (approximately seven generations), 1 mL of culture (~450,000 cells) was transferred into 150 mL of fresh medium. Every six weeks, 25 mL of each population were collected prior to transfer, pelleted and stored at −80 °C for DNA analysis. Every ~100 generations, DNA samples were extracted with the Qiagen DNeasy PowerBiofilm kit and then sent to the University of Pittsburgh Microbial Genome Sequencing Center for library construction and Illumina sequencing, as above. Sequence data have been deposited in the SRA under NCBI BioProject number ###########.

### Mutation detection

We used *breseq* v0.33.2 [64] to identify mutations and their frequencies in the ancestral and experimental populations with the strain CCMEE 5410 ancestral genome assembly as reference. FASTQ data were first quality-trimmed using *Trimmomatic* v0.39 (ILLUMINACLIP:TruSeq3-PE:2:30:10 HEADCROP:15 CROP:135 SLIDINGWINDOW:4:20 MINLEN:135; [58]). *breseq* analyses were performed in polymorphism mode with the default mutation frequency detection cut-off of 5%. For each candidate mutation, we used Fisher’s exact tests to test for biased strand representation and Kolmogorov-Smirnov tests to evaluate whether bases supporting a mutation had lower quality scores than those supporting the reference. We also confirmed candidate mutations by manually inspecting the alignments of reads to the reference genome.

### Phenotypic assays

After 400 generations of laboratory evolution, we assayed growth of the ancestral and evolved populations. Cells of the ancestral population were revived from a frozen stock stored at −80 °C. Cells of each population were rinsed with fresh medium and then used to inoculate triplicate flasks (each containing 150 mL of fresh FeMBG-11/HEPES medium) to a starting OD750 of ~0.015. Every 48 h, culture optical density at 750 nm (OD750) was measured with a Beckman Coulter DU 530 spectrophotometer (Indianapolis, IN). Generation times were estimated from the exponential growth phase of each culture.

Cell sizes of the ancestral and experimental populations were measured by imaging cells at 400X magnification with a Leica Model DME Microscope (Buffalo, NY). Images were then input into ImageJ 1.52q (National Institutes of Health) and 20 cells were measured after setting the appropriate pixel scale to obtain an average cell size for each culture.

To monitor cell aggregation, we modified the crystal violet adherence assay of Hernández-Prieto et al. [38]. Briefly, 2 mL of cell culture were inoculated in individual wells of 24 well-microplates at an OD750 about 0.13 (mid-exponential phase) at the start of the experiment. These immobile culture plates were grown for 10 days in cool white light at 30 °C. After incubation, the medium containing no adherent cells was decanted from each of the wells, and wells were rinsed gently with fresh FeMBG-11 media. To measure the number of adherent cells, each well was stained with 0.5 mL of 0.1 % crystal violet (CV) in ddH_2_O for approximately 1 hr. Once the CV solution was removed, wells were gently rinsed with ddH_2_0. The adherent cells were then resuspended in 2 mL 70 % ethanol for 15 min. Absorbance at 595 nm was used as a measurement of the number of cells adhered to the surface. Two technical replicates were performed for each biological replicate (*N* = 3).

## Acknowledgements

This work was supported by award NNA15BB04A from the National Aeronautics and Space Administration to S.R.M. S.R.M. thanks the Instituto Gulbenkian de Ciência for its support and hospitality during the analysis and writing of this project, and we thank Isabel Gordo, Massimo Amicone, Paulo Durão and Nelson Frazão for their insightful comments on an earlier version of the manuscript.

## Figure supplements, source data files and supplementary files

**Figure 1 – figure supplement 1.**
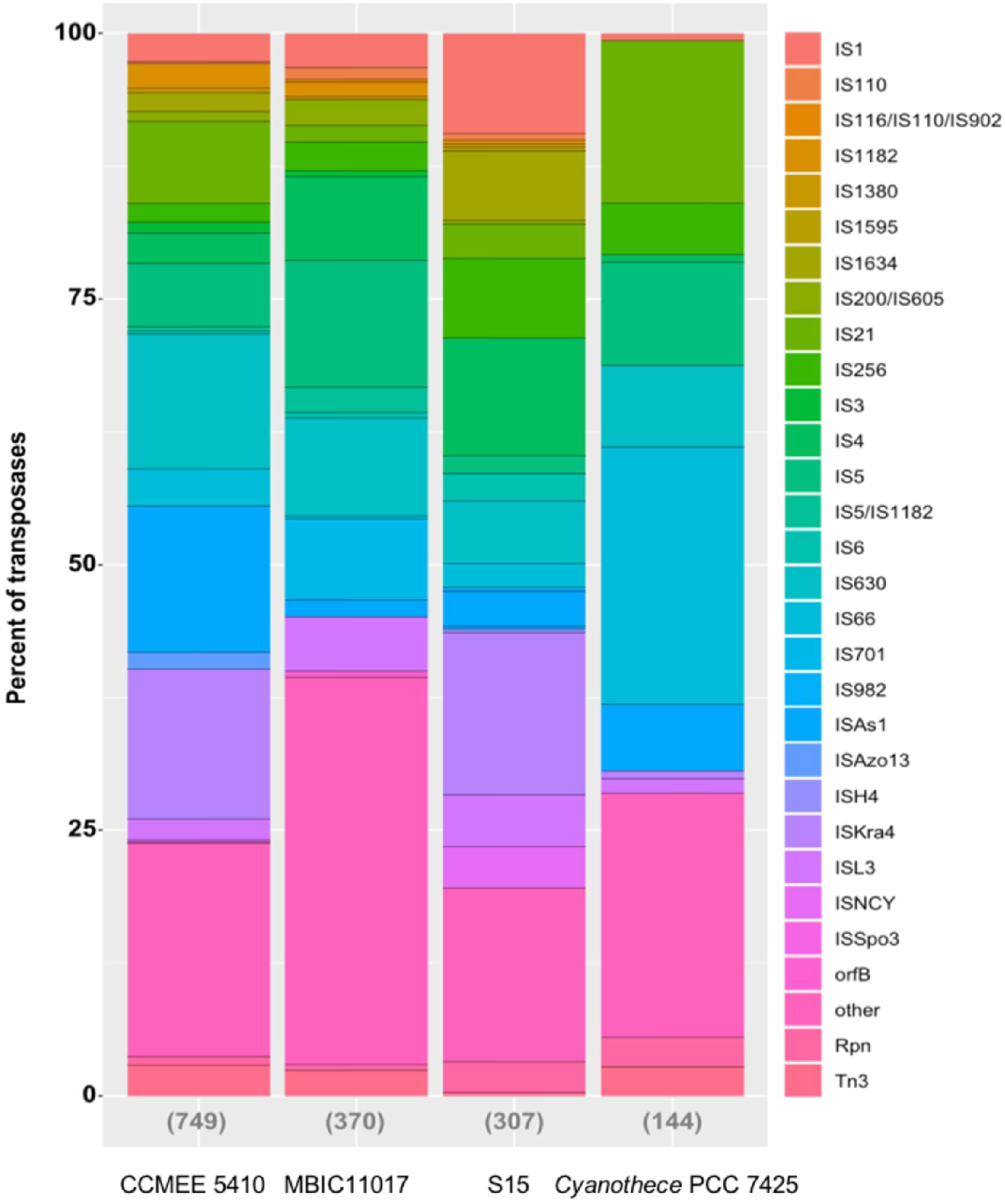
Relative frequencies of different IS families in *Acaryochloris* and *Cyanothece* genomes. The total number of transposase genes in each genome are indicated in parentheses.

**Figure 1 – figure supplement 2.**
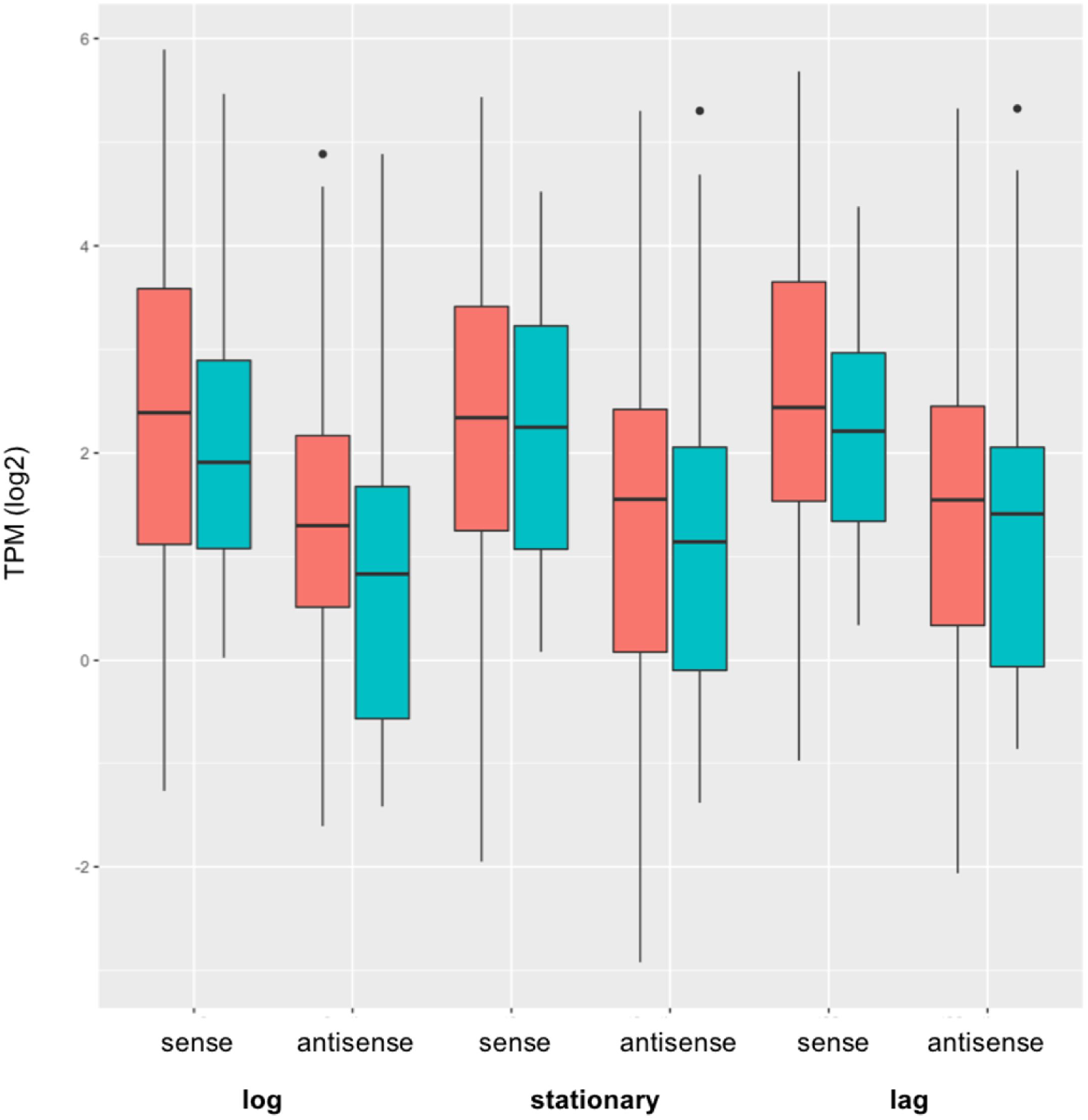
Log2 expression (TPM) of sense and antisense transcripts for high d*N*/d*S* (orange) and low d*N*/d*S* (blue) classes of IS elements in the *A. marina* strain CCMEE 5410 genome during log, stationary and lag phases of the population batch growth cycle. Sense transcripts from the high d*N*/d*S* class were significantly more highly expressed than the low d*N* /d*S* class in both log phase (*F*_1,159_ = 4.43, p = 0.037) and lag phase (*F*_1,159_ = 6.29, p = 0.013).

**Figure 1 – source data 1.** This file contains the data used in figure supplement 1. Distribution of IS element families in *Acaryochloris* and *Cyanothece* PCC 7425 genomes.

**Figure 1 – source data 2.** This file contains the expression data used in Figure 1 panel C.

**Figure 2 – figure supplement 1.**
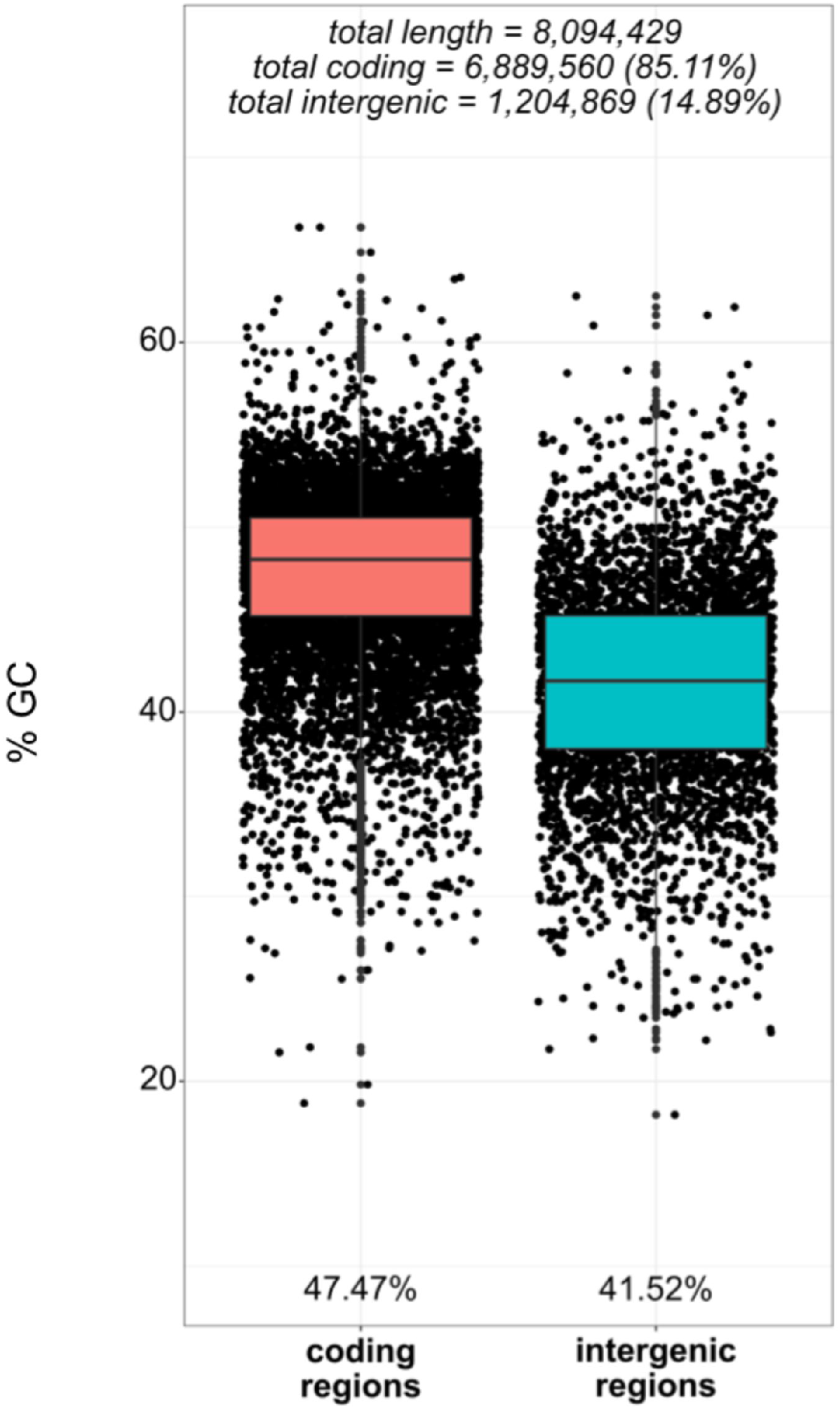
GC content of coding and intergenic regions of the *A. marina* strain CCMEE 5410 genome. Coding regions included all CDS, tRNA, rRNA, and tmRNA genes. Intergenic GC content was calculated only for those intergenic regions that are longer than 100bp.

**Figure 2 – figure supplement 2.**
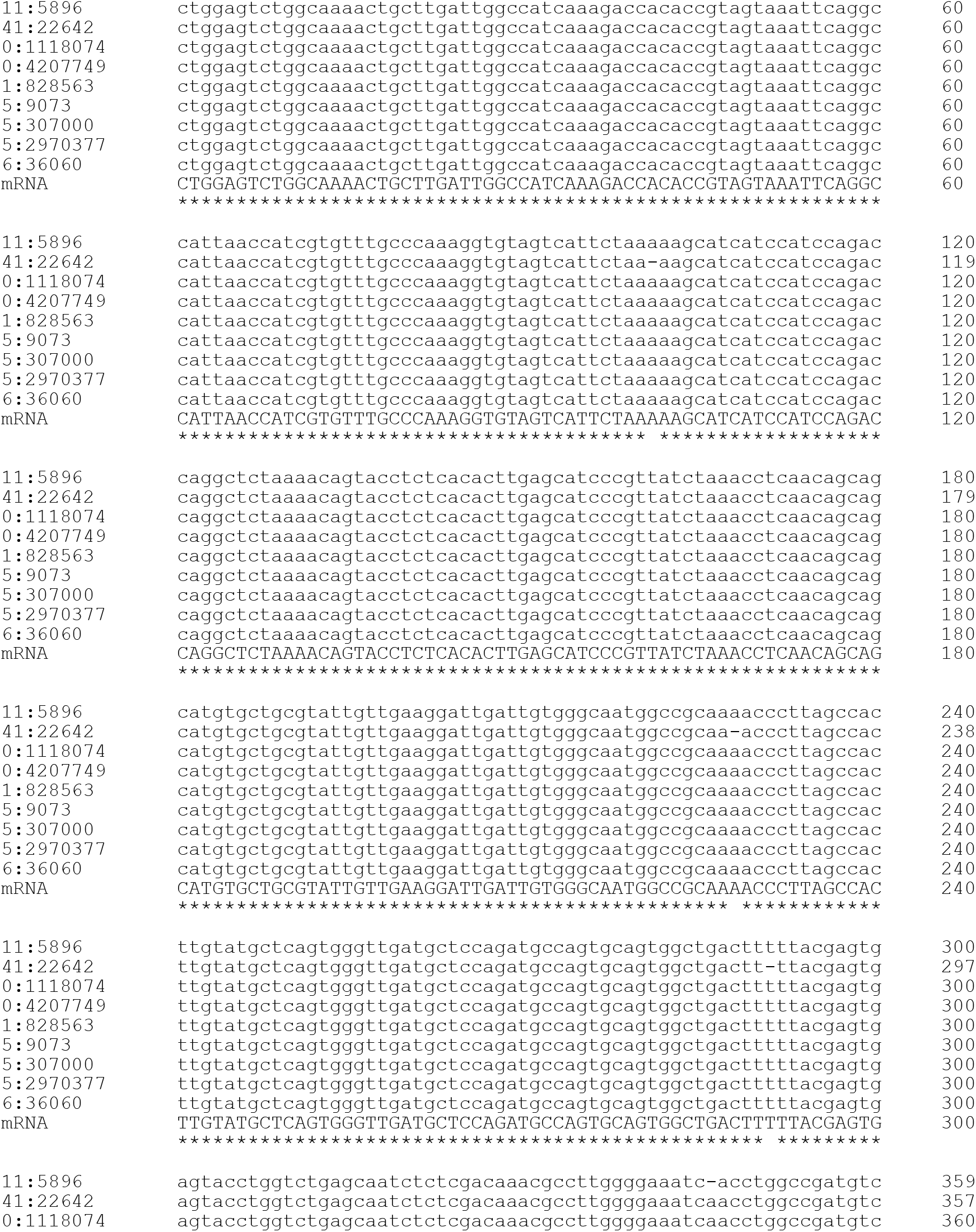
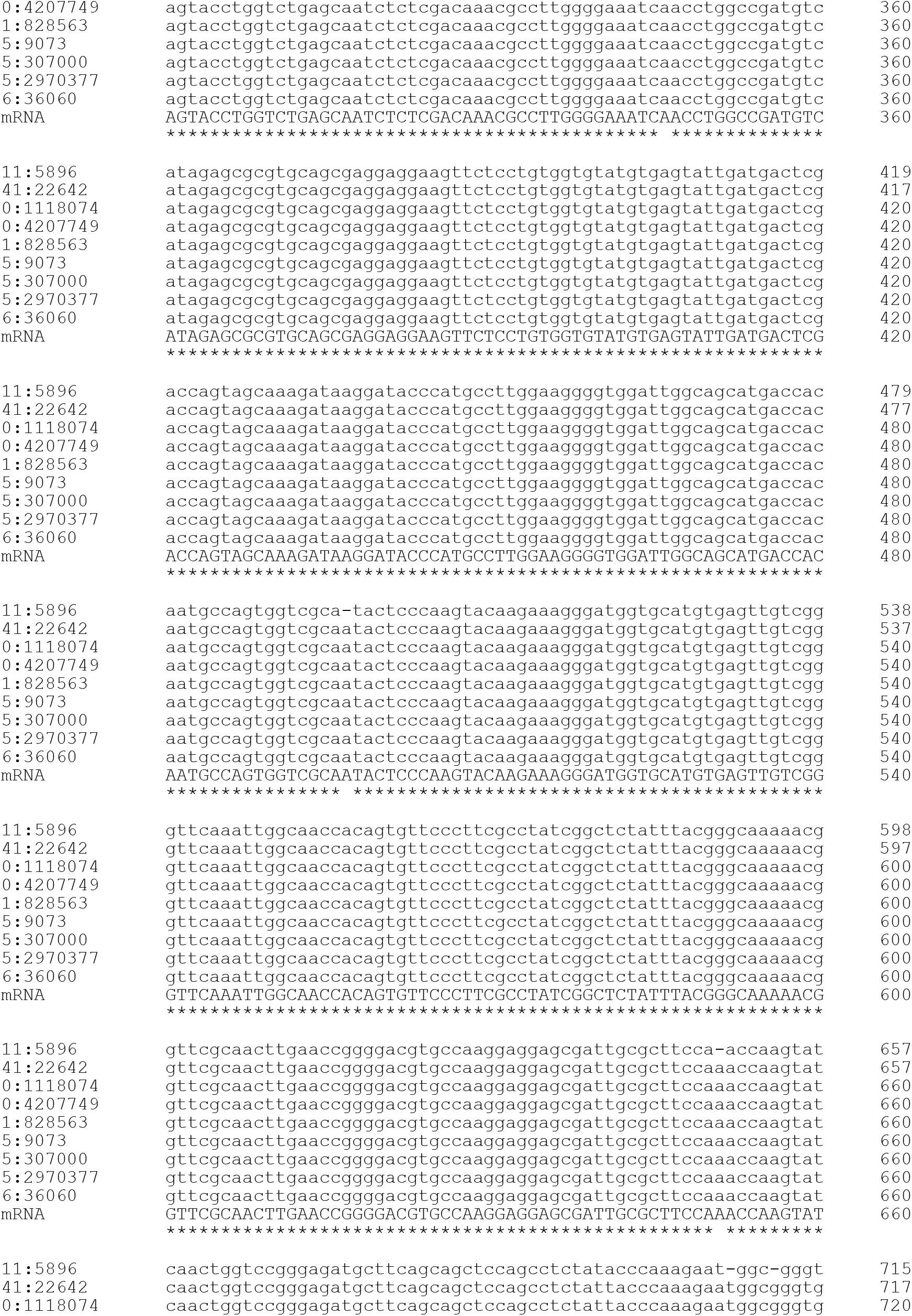
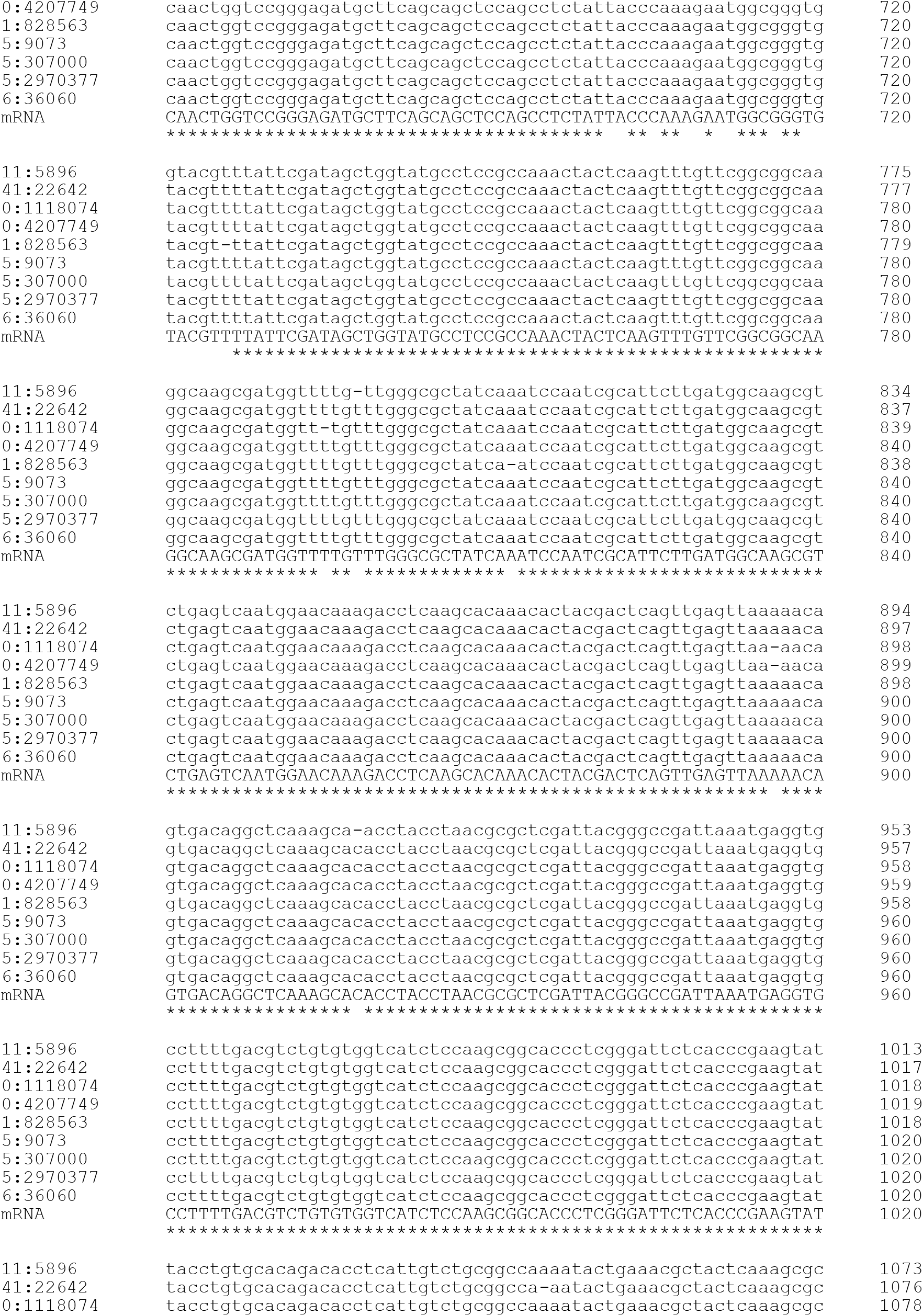
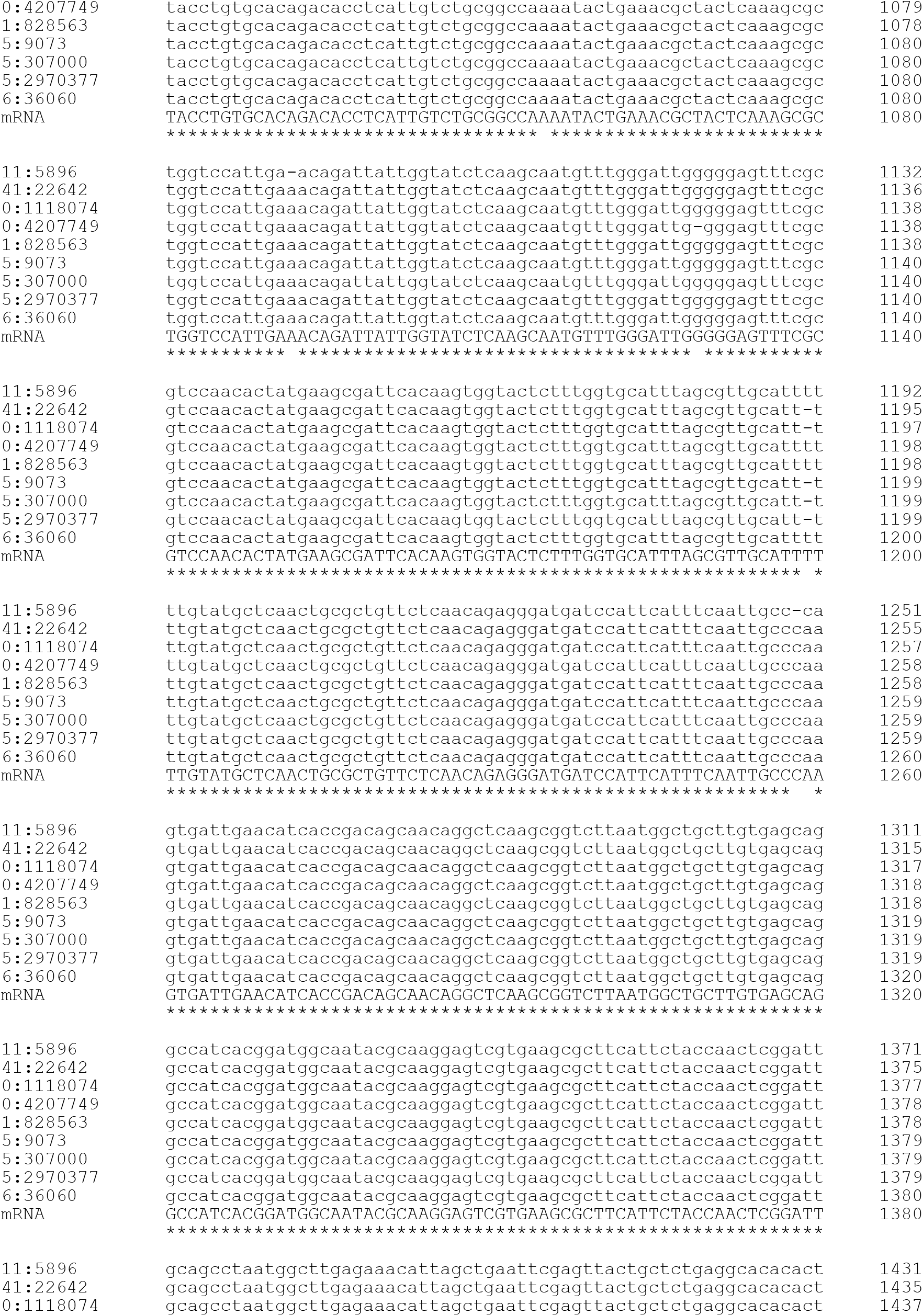
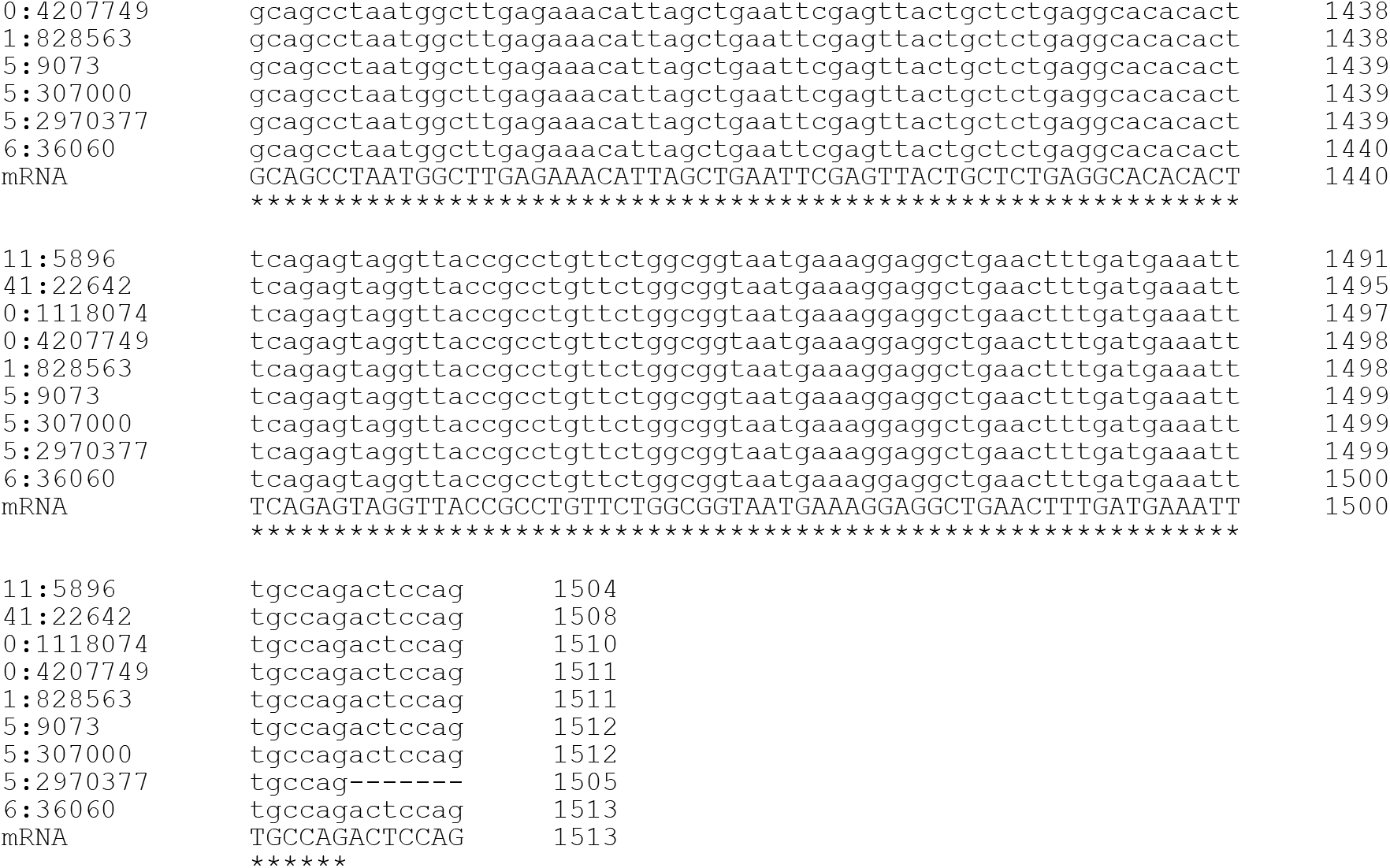
Nucleotide sequence alignment of the ISAm1 element reconstructed transcript and gene copies in the *A. marina* strain CCMEE 5410 genome (labels are genome coordinates). The transcript sequence is identical to the single complete copy of the element (6:36060).

**Figure 2 – source data 1.** This file contains the mutation data used in Figure 2 panel B.

**Figure 3 – figure supplement 1.**
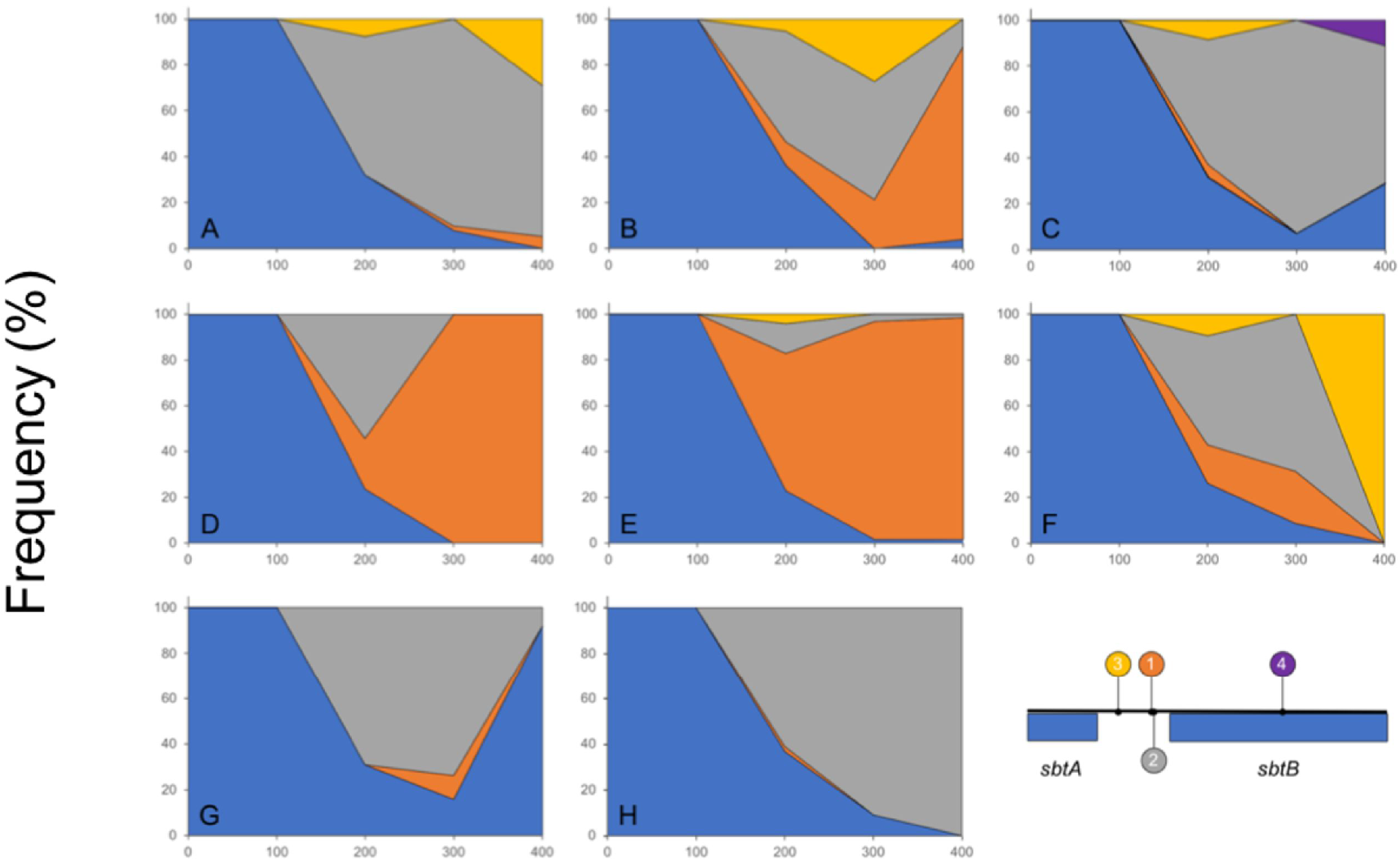
Relative frequencies of the ancestral Sbt allele (blue) and ISAm-1 insertion mediated mutations (see inset) in the eight populations during laboratory evolution. Inset: Location and frequencies of the four mutations in *sbtAB* detected during 400 generations of laboratory evolution. Shown is a 728 bp region of the CCMEE 5410 genome including the 3’ end of *sbtA*, intergenic DNA and *sbtB*.

**Figure 3 – source data 1.** This file contains the allele frequency data used in the fish plots.

**Supplementary file 1.** Summary of frameshifted transposase genes in *A. marina* genomes.

**Supplementary file 2.** Representative batch culture growth curve for *A. marina* CCMEE 5410 during laboratory evolution. Growth was monitored by the increase in optical density at 750 nm, which is proportional to cell density.

**Supplementary file 3.** Genome sequence coverage for laboratory evolved populations.

**Supplementary file 4.** Sense and anti-sense gene expression at different batch culture growth phases in the *A. marina* strain CCMEE 5410 ancestor for representative IS elements that contributed to laboratory evolution.

**Supplementary file 5.** Polymorphisms in the *A. marina* strain CCMEE 5410 ancestral population.

**Supplementary file 6.** Gene expression in the *A. marina* CCMEE 5410 ancestor under different growth conditions for mutated genes and select carbon-concentrating mechanism genes.

**Supplementary file 7.** Custom Python scripts used in this study.

